# Genome-wide binding of Multiple ankyrin repeats single KH domain reveals its role in maintenance of gene activation by trithorax group proteins in *Drosophila*

**DOI:** 10.1101/2022.03.15.484481

**Authors:** Ammad Shaukat, Muhammad Haider Farooq Khan, Jawad Akhtar, Mahnoor Hussain Bakhtiari, Muhammad Abdul Haseeb, Khalida Mazhar, Zain Umer, Muhammad Tariq

**Affiliations:** Epigenetics and Gene Regulation Laboratory, Department of Life Sciences, Syed Babar Ali School of Science and Engineering, Lahore University of Management Sciences, Lahore 54792, Pakistan

**Keywords:** Mask, ANKHD1, transcriptional cell memory, Trithorax, histone modifications, H3K27ac

## Abstract

The Trithorax group (trxG) proteins counteract repressive effect of Polycomb group (PcG) complexes and maintain transcriptional memory of active states of key developmental genes. Although, chromatin structure and modifications appear to play a fundamental role in this process, it is not clear how trxG prevents PcG-silencing and heritably maintain an active gene expression state. Here, we report a hitherto unknown role of *Drosophila* Multiple ankyrin repeats single KH domain (Mask), which emerged as one of the candidate trxG genes in our reverse genetic screen. The genome-wide binding profile of Mask correlates with known trxG binding sites across *Drosophila* genome. In particular, association of Mask at chromatin overlaps with CBP and H3K27ac, which are known hallmarks of actively transcribed genes by trxG. Importantly, Mask predominantly associates with actively transcribed genes in *Drosophila*. Depletion of Mask not only results in downregulation of trxG targets but also correlates with drastic reduction in H3K27ac levels and an increased H3K27me3 levels. The fact that MASK positively regulates H3K27ac levels in flies was also found to be conserved in human cells. Finally, strong suppression of *Pc* mutant phenotype by mutation in *mask* provides physiological relevance that Mask contributes to the anti-silencing effect of trxG, maintaining expression of key developmental genes. Since Mask is a downstream effector of multiple cell signaling pathways, we propose that Mask may connect cell signaling with chromatin mediated epigenetic cell memory governed by trxG.

## Introduction

In multicellular eukaryotes, differential gene expression profiles established during the patterning processes lead to specialization of cells during early embryonic development. In order to grow and maintain a specialized state, the particular expression profiles of genes need to be transmitted to daughter cells in all cell lineages. The Polycomb (PcG) and Trithorax (trxG) group proteins maintain the transcriptional states of important developmental regulators like the *Hox* genes which ensure that determined cell fates are faithfully propagated during cell proliferation [1–5]. Both the PcG and trxG proteins act in large multi-protein complexes and modify the local properties of chromatin to maintain transcriptional states of repression and activation of their target genes, respectively [6].

The association of PcG and trxG proteins with chromatin is evolutionarily conserved and defects in PcG/trxG system lead to severe morphological abnormalities during development [7,8]. Although, chromatin structure and modifications appear to be important, genome organization has also emerged as an important factor in transcriptional regulation by PcG/trxG [9].

In *Drosophila,* PcG and trxG proteins are known to exert their functions by binding to specific chromosomal elements known as *PREs* (Polycomb Response Elements) [10,11]. Gene silencing by the Polycomb Repression Complex 1 (PRC1) and Polycomb Repression Complex 2 (PRC2) employs an intricate interplay of different components that regulate the target genes in a spatio-temporal manner during development. Importantly, histone H3 lysine 27 trimethylation (H3K27me3), histone H2A lysine 118 ubiquitination (H2AK118ub) and stalled RNA polymerase II at gene promoters are considered hallmarks of PcG-silencing. Although the trxG proteins are known to directly counteract repression by the PcG, the maintenance of gene activation by trxG is seen as a constant battle against the default state of silencing by PcG. Details of the molecular mechanism employed by trxG to prevent PcG-silencing still remain elusive [12,13]. The trxG proteins display a wide range of functional diversity which includes covalent modifications of histones, chromatin remodeling and general transcriptional processes. It can be envisaged that the function and composition of trxG complexes is modulated in various cell types at specific target genes [14]. Due to the intricate nature of gene activation by trxG, it is important to identify the as yet unknown accessory proteins that may interact with the trxG in maintenance of gene activation.

Our laboratory employed an *ex-vivo* genome-wide RNAi approach to identify novel trxG factors involved in maintaining gene activity. More than 200 different genes were identified as potential trxG regulators. Mask, a large scaffold protein, was also among the potential trxG candidate genes [15]. Initially, Mask was discovered in *Drosophila* as a regulator of Epidermal Growth Factor (EGF) signaling. It was reported to play an important role in photoreceptor differentiation, cell survival and proliferation [16]. *Drosophila* Mask is a large nuclear as well as cytoplasmic protein which contains 4001 amino acids harboring different functional domains. Notably, Mask has been linked to different cell signaling pathways and various cellular processes in proliferating [17–19] and non-proliferating cells [20]. Besides EGF signaling, Mask is known to act downstream of Hippo signaling pathway as a co-transcription factor, required for organ size regulation during development [17–19].

Here we report a previously unknown role of Mask in the maintenance of gene activation by trxG in *Drosophila* and humans. We discovered that the genome-wide binding profile of Mask massively overlaps with Trx protein binding sites in *Drosophila.* Importantly, Mask associates with known *PREs* within Antennapedia complex (ANT-C) and Bithorax complex (BX-C) regions, which are canonical targets of trxG/PcG [21]. Mask is also enriched at histone H3 lysine 27 acetylation (H3K27ac) marked chromatin, a histone mark deposited by CREB binding protein (CBP) and a hallmark of trxG mediated gene activation [22]. In corroboration, our analysis also revealed an overwhelming correlation between Mask and CBP binding at active chromatin. Depletion of Mask in *Drosophila* and mammalian cells resulted in a drastic reduction in global H3K27ac levels and an increase in H3K27me3 levels. The co-occupancy of Mask and CBP on chromatin further substantiates the effect of Mask on H3K27ac levels. Finally, strong suppression of *Pc* associated extra sex comb phenotype by *mask* mutant, analogous to the trxG effect, supports the notion that Mask positively regulates maintenance of active chromatin states by trxG. Further molecular and biochemical investigation will decipher mechanistic details of how MASK influences CBP mediated H3K27ac and consequent maintenance of gene activation. Answers to these questions may reveal the missing link between cell signaling, trxG and cell fate maintenance.

## Results

### Mask regulates trxG target genes and associates with polytene chromosomes

Since Mask appeared as a potential trxG gene in an *ex-vivo* genome-wide RNAi screen based on luciferase reporter in *Drosophila* [15], we first validated whether Mask is able to regulate expression of endogenous trxG target genes. To this end, we knocked down *mask* and analyzed the expression of, *pannier (pnr), pointed (pnt)* and *pipsqueak (psq),* known targets of trxG [22]. Depletion of Mask resulted in downregulation of trxG targets similar to the effect of *trx* knockdown in these cells (Figure 1A). This highlights a potential role of Mask in maintenance of gene activation by trxG in *Drosophila.* Since trxG/PcG proteins influence gene expression through their association with chromatin [5,23], it was investigated if Mask also binds to chromatin. To determine the chromatin association of Mask, we isolated salivary glands from third instar larvae of *w^1118^* flies and stained polytene chromosomes with anti-Mask antibody. Notably, Mask was found to localize on inter-band regions on polytene chromosomes, which indicates its predominant binding on potentially actively transcribed regions (Figure 1B-D).

**Figure 1:**
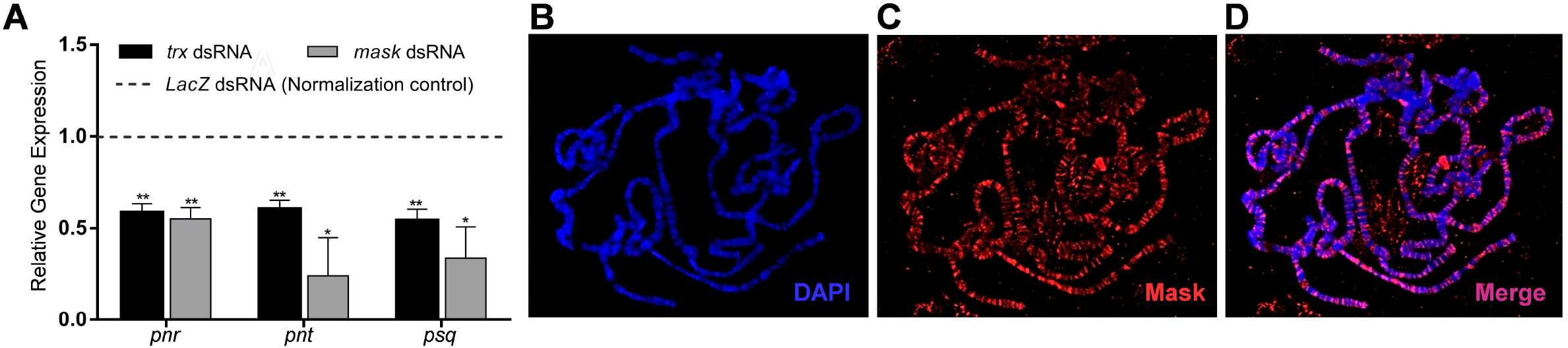
Mask regulates trxG targets and binds to inter-band regions on polytene chromosomes. **(A)** Relative gene expression of *pnr, pnt* and *psq* measured by qRT-PCR, after knockdown of *mask* in D.Mel-2 cells. Cells treated with *LacZ* dsRNA served as normalization control (dashed line) while Trx depleted cells were used as positive control. Depletion of Mask results in downregulation of *pnr, pnt* and *psq*, similar to the *trx* knockdown. Experiment was repeated three times and independent t-tests were performed for each target gene (*p ≤ 0.05, **p ≤ 0.01). **(B-D)** Immunostaining of polytene chromosomes from 3^rd^ instar larvae of *w^1118^* with DAPI **(B)** and anti-Mask antibody **(C)** reveals binding of Mask mainly on interband regions in polytene chromosomes.

### Genome-wide binding profile of Mask overlaps with Trx binding at active genes

After observing binding of Mask on polytene chromosomes, the genome-wide binding profile of Mask on chromatin was investigated. To this end, chromatin immunoprecipitation sequencing (ChIP-seq) from *Drosophila* cells was performed using anti-Mask antibody. Mask was found to bind 2848 regions and 69% of these binding sites were in promoters, whereas few sites covered introns and distal intergenic regions (Figure 2A). The genome-wide binding profile of Mask further indicated that Mask occupies chromatin regions mainly near ±2 kb of TSS (Transcription Start Site) of genes (Figure 2B, C). Since Trx protein is proteolytically cleaved into Trx-N and Trx-C proteins and both bind to chromatin, heat maps of previously known genome-wide binding profiles of Trx-C, Trx-N, H3K27ac, H3K27me3 [24] and Pc [25] were plotted within ±10 kb of TSSs of genes bound by Mask to analyze overlay with Mask binding profile (Figure 2D). This analysis revealed a significant overlap of Mask-bound sites with previously reported Trx-C, Trx-N and Pc ChIP-seq profiles. In addition, a strong correlation of Mask binding with H3K27ac marked chromatin was observed. In contrast, very less overlap of Mask binding profile was observed with H3K27me3 associated chromatin (Figure 2D). These results clearly indicate a potential role of Mask in gene activation that was validated by comparison of Mask binding sites with previously reported actively transcribed genes in *Drosophila* cells [26] (Figure 2E). Additionally, Mask enrichment at known enhancers, such as *Ptx1* and *Optix* [24], also suggests its potential role in gene activation (Supplementary Figure 1A, B). A comparison of short motifs enriched in ChIP-seq data of Mask and previously reported ChIP-seq data of Trx revealed similarity in top-scoring motifs. Most importantly, GAGA motif, a known GAGA factor (GAF) binding site and GCCAT motif, a known PHO binding motif were enriched in case of both Mask and Trx bound chromatin regions [27] (Figure 2F). An overwhelming overlap of Mask binding sites with previously reported genome-wide binding profiles of Trx and Pc within BX-C (Figure 3A) and ANT-C (Figure 3B) was observed where Mask was found to occupy majority of all known *PREs* [21,28]. Analysis of Mask binding sites at cell signaling genes revealed that Mask associates with thirty-three different genes belonging to Wnt signaling pathway, including *en* (Supplementary Table 1). The Trx and Pc binding sites at *PREs* near *en* promoter [29–31] also show a strong overlap between Mask, Trx and Pc (Figure 3C).

**Figure 2:**
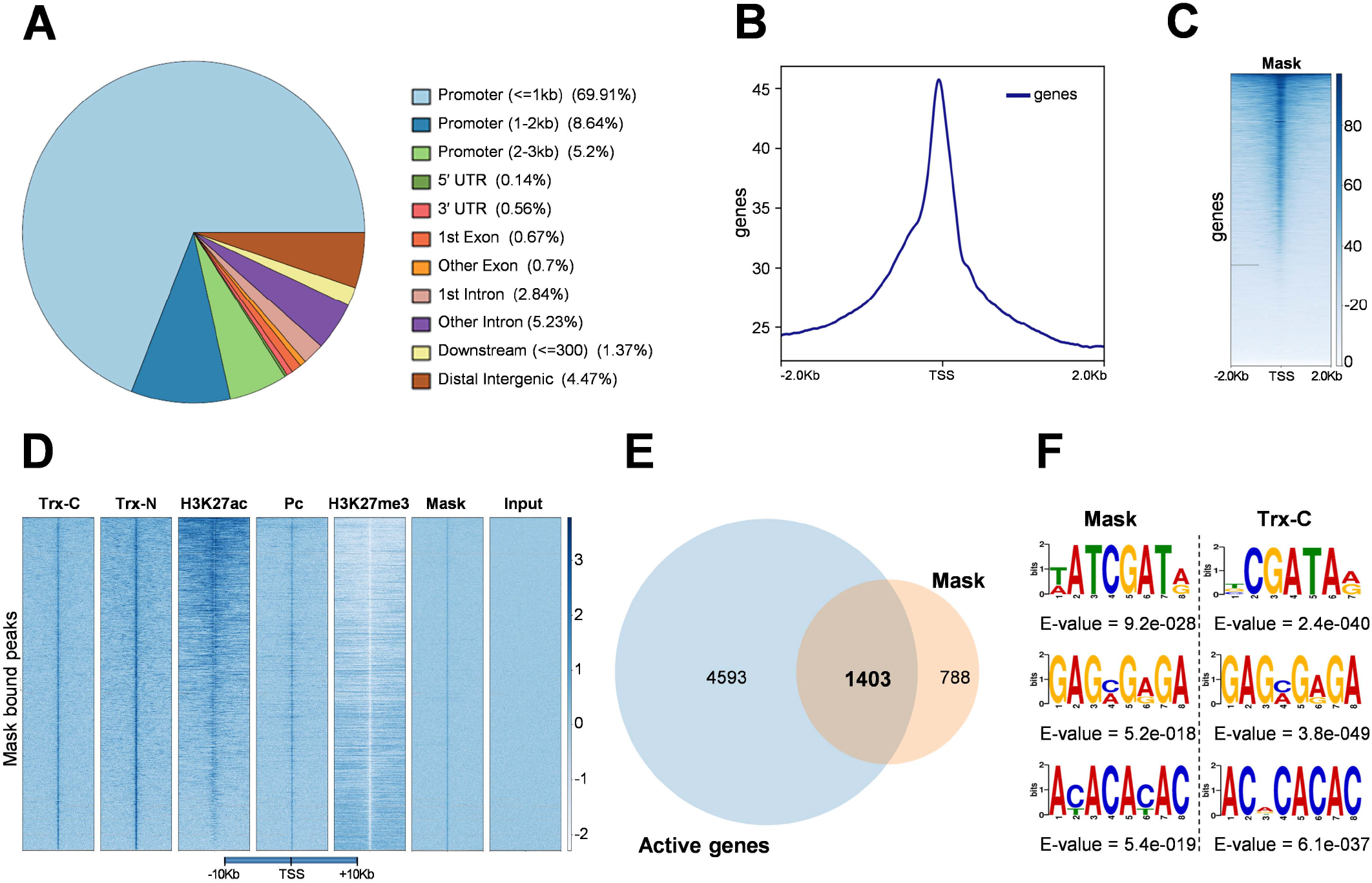
Genome-wide binding profile of Mask highlights its role in gene activation. **(A)** ChIP-seq data of Mask shows that Mask binding sites are mainly promoter regions across fly genome. Besides promoter regions, Mask was observed at very few UTR’s, introns, downstream and distal intergenic regions. **(B)** Analysis of Mask enrichment across ±2Kb of TSS highlights its association predominantly at TSS of target genes. **(C)** Heat map depicting Mask enrichment across ±2Kb of TSS, which highlights its predominant binding at TSS. **(D)** Heat maps showing enrichment of Trx-C, Trx-N, H3K27ac, Pc and H3K27me3 within ±10 Kb of TSSs of genes bound by Mask. Strong enrichment of Trx-C, Trx-N, H3K27ac and Pc was observed on Mask bound regions at TSS. However, relatively little to no overlap between Mask binding and H3K27me3 was observed. **(E)** Venn diagram representing Mask bound genes superimposed with the list of actively transcribed genes in S2 cells. Most of the Mask associated genes were found to be transcriptionally active. **(F)** Top scoring short motifs of DNA enriched in Mask bound chromatin, generated from ChIP-seq data using MEME-ChIP database, show very high similarity with the DNA motifs in Trx ChIP-seq data. The frequency with which each nucleotide occurs is determined by the proportion of the total height of each letter. Motifs are presented with their respective E-values.

**Figure 3:**
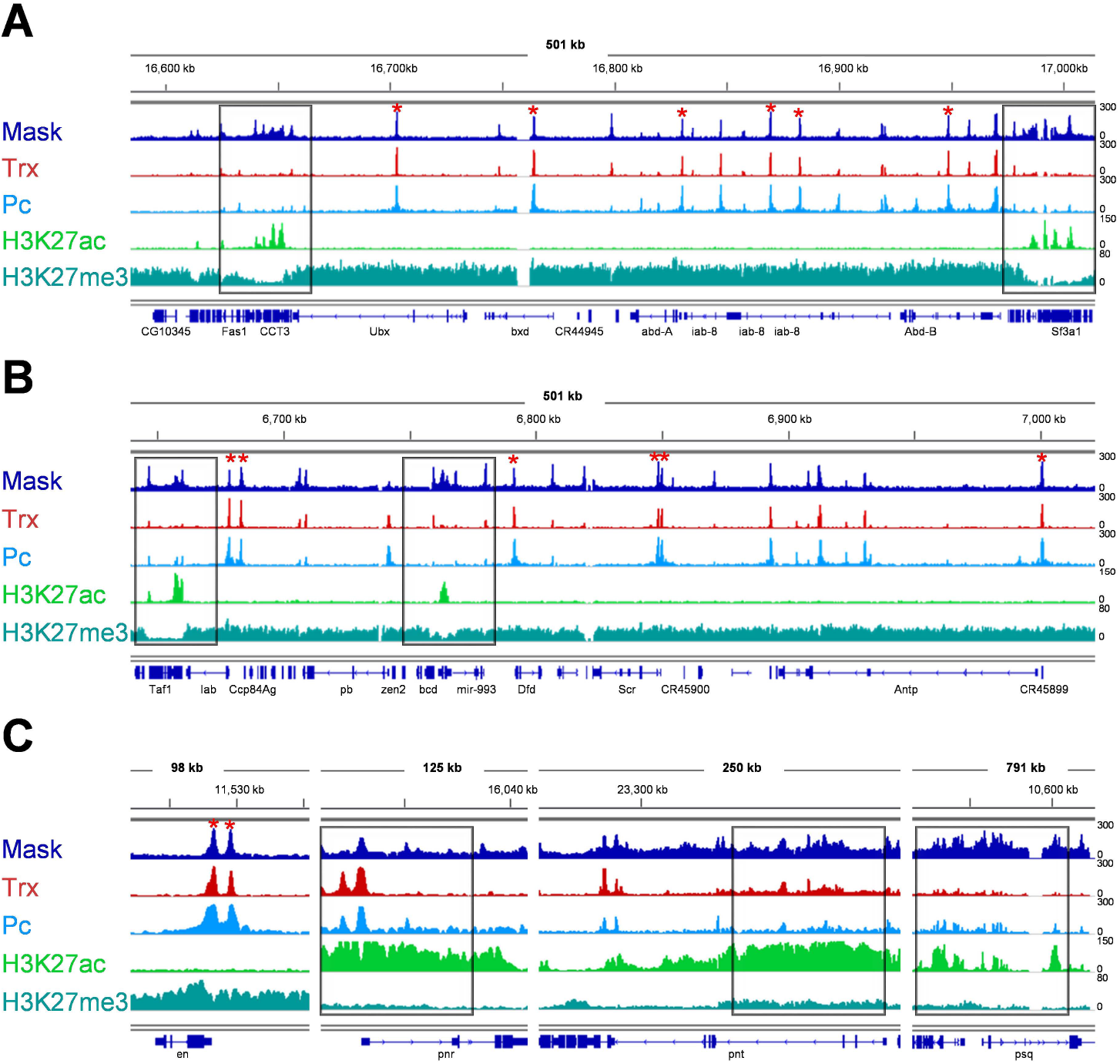
Mask associates with known Trx and Pc binding sites in homeotic and non-homeotic genes. **(A-C)** Integrated genome browser view (IGV) of ChIP-seq data of Mask in comparison with Trx, Pc, H3K27ac and H3K27me3 at various loci known to be bound by Trx and Pc. Mask occupies several known binding sites of Pc and Trx in Bithorax complex (BX-C) **(A)** and Antennapedia complex (ANT-C) **(B).** All the peaks marked with red asterisk represent known *PREs* in BX-C and ANT-C [60] **(A-B). (C)** Mask occupies known Trx and Pc binding sites in *engrailed (en)* locus, which corresponds to known *PREs* (marked with asterisk) in *en* gene. Mask was also highly enriched at regions where high levels of H3K27ac are present (regions marked with rectangle boxes shown in **A**, **B** and **C**), including *pnr, pnt* and *psq* loci.

### Mask overlaps with CBP binding on chromatin and positively regulates H3K27ac

A strong correlation was observed between Mask and H3K27ac across the genome as exemplified by the *CCT3* locus adjacent to BX-C (Figure 3A, Supplementary Figure 1C). Additionally, Mask binding at *pnr, pnt* and *psq* loci was investigated (Figure 3C), the genes which are downregulated after depletion of either Mask or Trx (Figure 1A). It was observed that Mask was strongly enriched on these actively transcribed genes, which correlates with higher enrichment of H3K27ac. To dissect the molecular interaction between Mask and H3K27ac, it was investigated if Mask shares chromatin binding sites with CBP, an enzyme that catalyzes H3K27ac. To this end, ChIP-seq data of Mask was compared with previously published CBP binding profile [32] from *Drosophila* S2 cells. This analysis revealed a strong overlap in Mask, CBP and H3K27ac genome wide profiles in *Drosophila* (Figure 4A). Heat map representing genome-wide binding profile of CBP covering ±10 kb of TSSs of genes bound by Mask reveals strong correlation in their binding pattern (Figure 4B). Further analysis revealed that both Mask and CBP shares 2506 binding sites on chromatin (Figure 4C, Supplementary Table 1). Interestingly, the genomic distribution of Mask (Figure 2A) and CBP [26] in terms of their binding at promoter regions, gene body, introns and intergenic regions also reveal a striking similarity. Together, this data highlights that Mask binding pattern substantially overlaps with the binding profile of CBP. These findings prompted us to investigate if depletion of Mask affects CBP mediated H3K27ac levels in *Drosophila.* To this end, *mask* was knocked down using RNAi in *Drosophila* S2 cells and global levels of H3K27ac were analyzed on a Western blot. As compared to control cells, total H3K27ac levels were found to be strongly diminished in Mask depleted cells (Figure 4D). Importantly, increased H3K27me3 levels were also observed after Mask depletion (Figure 4D). Such a drastic effect of impaired Mask function on H3K27ac and H3K27me3 levels encouraged us to investigate conservation of this interaction in mammalian cells. To this end, we generated *ANKHD1* (mammalian homologue of *Drosophila* Mask) knockout (KO) cells using CRISPR/CAS system in HEK293T cells. As compared to control cells, strongly diminished H3K27ac levels were observed in *ANKHD1* KO cells (Figure 4E). Moreover, enhanced levels of H3K27me3 were also observed in *ANKHD1* KO cells, which corroborates the effect of Mask depletion in *Drosophila* cells. Importantly, elevated levels of EZH2 protein, a PRC2 component that catalyzes H3K27me3 [33], were also observed in *ANKHD1* KO cells as compared to normal cells (Figure 4E). All these results correlate with downregulation of highly acetylated genes after depletion of Mask and provide clues towards mechanistic interactions of Mask with CBP (Figure 1A, Figure 3C). Similarity in Mask and CBP chromatin binding patterns and effect of Mask on H3K27ac levels in flies and mammalian cells illustrates that Mask positively regulates H3K27ac (Figure 4D, E) and it is required for maintenance of gene activation by trxG.

**Figure 4:**
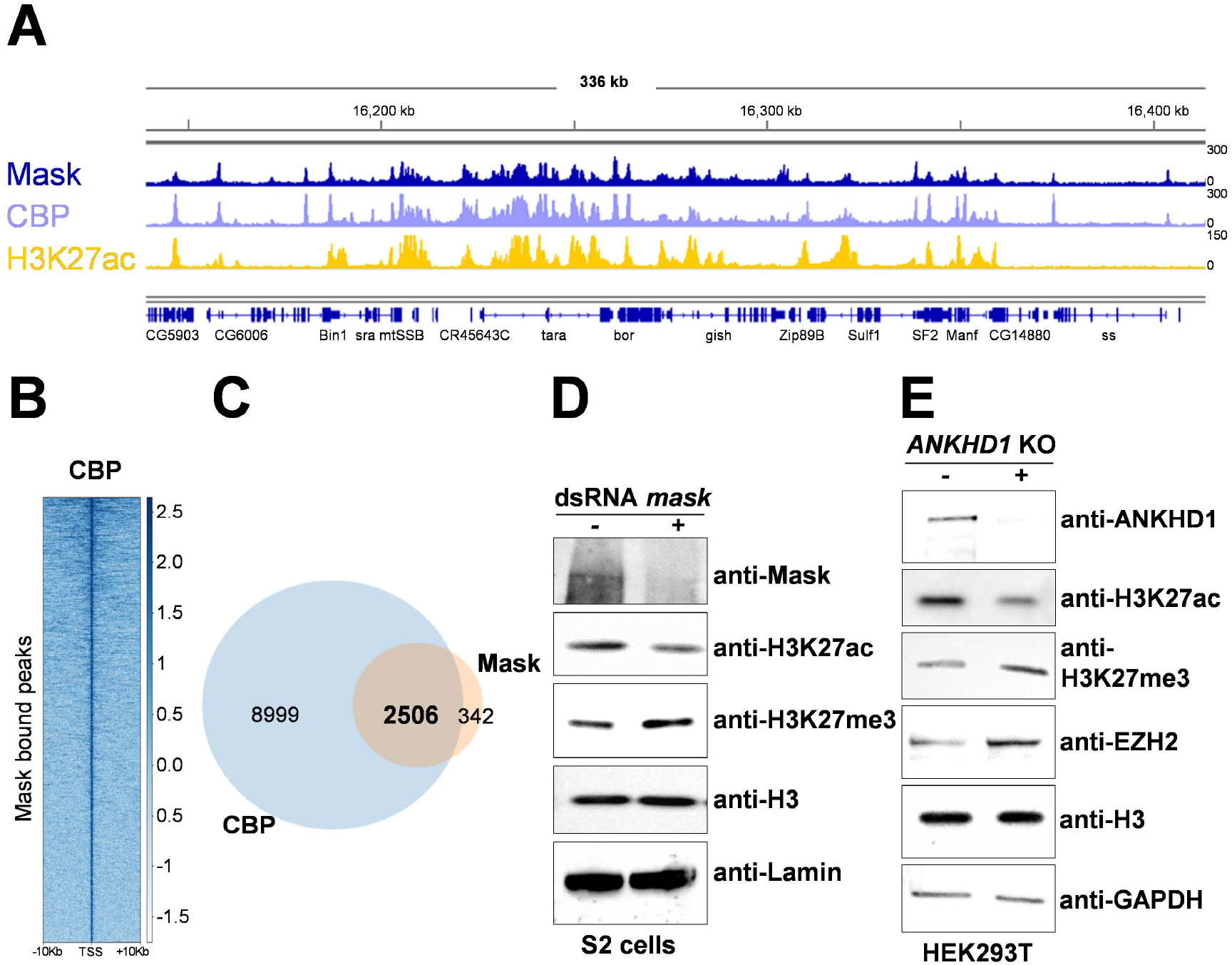
Genome-wide binding of Mask overlaps with CBP binding sites and Mask positively regulates H3K27ac levels. **(A)** Integrated genome browser (IGV) view of the ChIP-seq data shows striking overlap between genome-wide patterns of Mask, CBP and H3K27ac. **(B)** Heat map depicting enrichment of CBP within ±10 Kb of TSSs of genes bound by Mask reveals a strong overlap between Mask and CBP occupancy. **(C)** Venn diagram represents substantial overlap between Mask and CBP bound regions on chromatin. Both Mask and CBP co-occupy 2513 peaks. **(D)** Western blot analysis of whole cell lysates from *Drosophila* D.Mel-2 cells treated with dsRNA against *mask* (+) show depletion of MASK as compared to the cells treated with *LacZ* (-) dsRNA which served as a negative control. MASK depleted cells exhibit decreased H3K27ac levels and increase in H3K27me3 levels. Total levels of histone H3 and Lamin were used as a loading control. **(E)** Western blot analysis of HEK293T cells stably transfected with *ANKHD1* CRISPR knockout (KO) plasmids (+), exhibit drastically reduced H3K27ac levels as compared to untreated cells. H3K27me3 and EZH2 levels also increased in *ANKHD1* KO cells as compared to control (-). Total levels of histone H3 and GAPDH served as loading control in immunoblot.

### Mask exhibits trxG like behavior in *Drosophila*

Since Mask was found to bind at multiple sites within homeotic genes in BX-C and ANT-C (Figure 3A, B) in *Drosophila* S2 cells, we questioned if Mask association at homeotic genes has any physiological relevance during fly development. To this end, we crossed a *mask* mutant *(mask^EY09448^)* with two different *Pc* mutants *(Pc^1^, Pc^XL5^*) to observe effect of *mask* mutation on extra sex comb phenotype in *Pc* mutants. Normally, *Pc* heterozygous mutant flies show extra sex comb phenotype due to de-repression of *Sex comb reduced (Scr)* gene in second and third pair of legs [34]. In the *Pc* heterozygous mutant background, *mask^EY09448^* strongly suppressed the extra sex comb phenotype, indicating loss of *Scr* expression (Figure 5A, Supplementary Figure 2). Strong suppression of extra sex comb phenotype by *mask^EY0944^* suggests a trxG-like behavior which correlates with Mask binding on *Scr* locus in ChIP-seq data (Figure 3B). Since *mask* mutant exhibits trxG like behavior, it was investigated if *mask* genetically interacts with *trithorax (trx).* The genetic interaction between *mask* and *trx* was analyzed by crossing *mask* mutant *(mask^EY09448^)* with *trx* mutant (*trx^1^*) flies. The *trx* heterozygous mutant male flies show loss of pigmentation on the fifth abdominal segment (A5) due to decreased *Abdominal-B (Abd-B)* expression that controls dark pigmentation in male posterior body segments [35,36]. As compared to heterozygous *trx^1^* mutant males used as a control, *mask^EY09448^/trx^1^* trans-heterozygote male flies showed a significantly higher percentage of A5 to A4 transformation (Figure 5B). To further confirm this physiological evidence at the molecular level, we investigated if depletion of Mask in flies affects *en* gene expression, a known trxG/PcG target gene [29–31]. To this end, we crossed *UAS-mask^RNAi^* flies with a *GAL4* driver line (*en-GAL4,UAS-GFP*) to knockdown *mask* and observe its effect on *en* expression in the imaginal discs isolated from the third instar larvae. A cross between *en-GAL4,UAS-GFP* and *w^1118^* served as control. The control flies expressing *en-GAL4,UAS-GFP* alone show expression of GFP that overlaps with en staining in posterior compartment of imaginal discs where *en* is normally expressed [37] (Figure 5C). As compared to the control flies, expression of *UAS-mask^RNAi^* driven by *en-GAL4,UAS-GFP* driver resulted in drastic reduction in en staining with a concomitant reduction in GFP expression. Since GAL4 was under *en* enhancer, the decrease in both endogenous *en* expression and *en-GAL4* driven GFP, upon *mask* knockdown can be explained (Figure 5C). These findings were further confirmed by crossing *UAS-mask^RNAi^* with *GawB-GAL4* driver, which is normally expressed in pouch region of wing, leg and haltere imaginal discs (Supplementary Figure 3A). As compared to control flies that expressed *GawB-GAL4* driver alone, a drastic reduction in *en* expression was observed as a consequence of *mask* knockdown in progeny of flies expressing *mask^RNAi^* driven by *GawB-GAL4* driver (Supplementary Figure 3B). It is pertinent to mention that genome-wide binding profile of Mask revealed its association with known trxG/PcG binding sites within *en PRE* (Figure 3C), which further supports the drastic reduction of *en* expression after depletion of Mask in flies. Together, these results suggest that Mask is required for maintenance of gene activation of key developmental genes by trxG.

**Figure 5:**
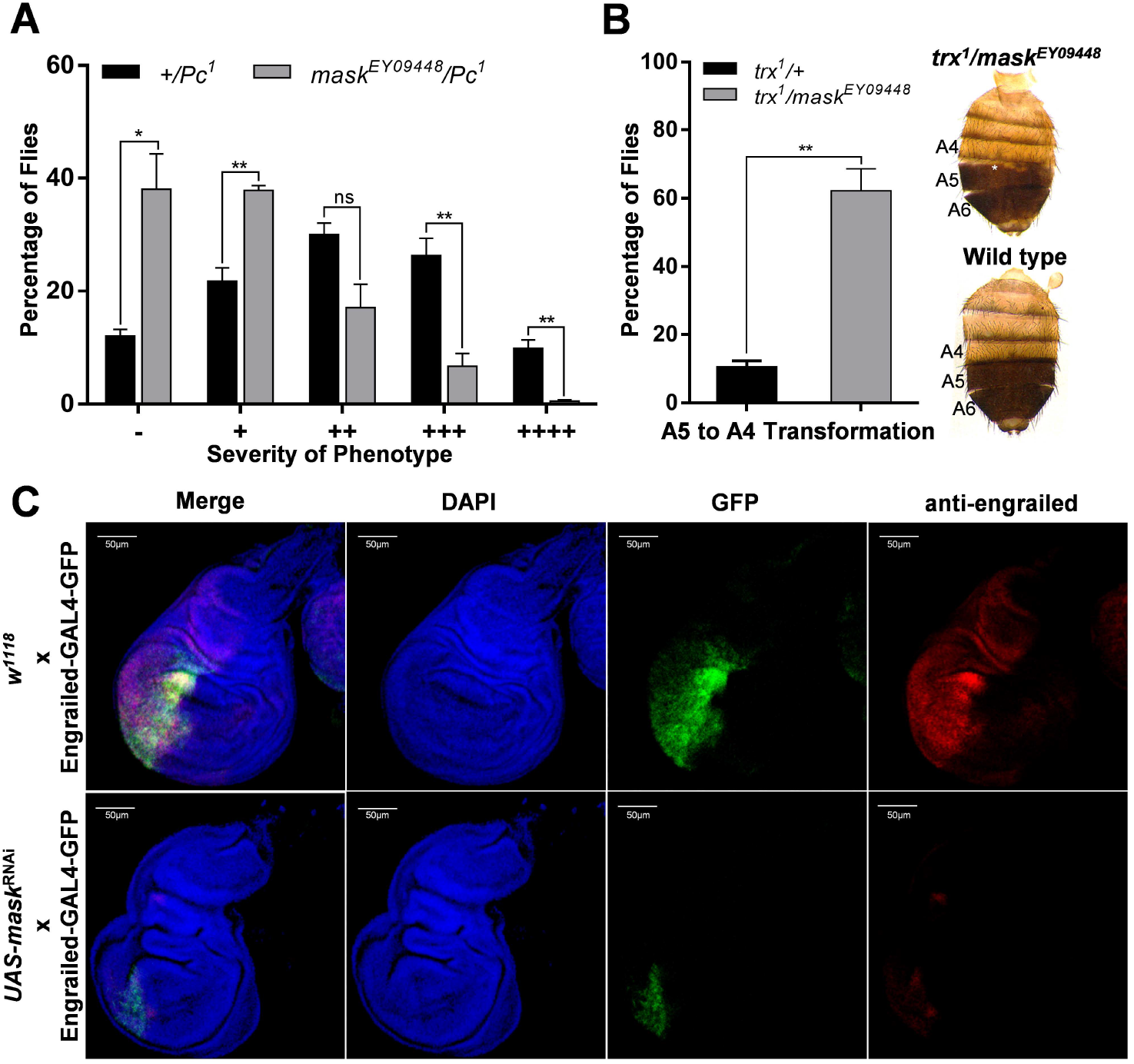
Mask genetically interacts with Pc/trxG system and exhibits trxG like behavior regulating trxG target genes in *Drosophila*. **(A)** Mutant of *mask (mask^EY09448^)* was crossed with *Pc* mutant (*Pc^1^*) and males in the progeny with *mask^EY09448^/Pc^1^* genotype were scored for extra sex comb phenotype. Males from *Pc^1^* flies crossed with *w^1118^* served as control. *Pc^1^*/+ exhibited a strong extra sex comb phenotype in contrast to *mask^EY09448^/Pc^1^*. Total 200 male flies of the desired genotype from progeny of each cross were analyzed and based on the presence of extra sex combs on 2^nd^ and 3^rd^ pair of legs, flies were categorized according to the severity of phenotype. These categories are: –, no extra sex combs; +, 1–2 hairs on 2nd leg; ++, more than three hairs on 2nd leg; +++, more than 3 hairs on 2nd leg and 1–2 hairs on 3rd leg; ++++, strong sex combs on both 2nd and 3rd pairs of legs as described previously [57]. The experiment was repeated three times independently and individual t-test was used to compare each phenotype category. **(B)** *mask* mutant, *mask^EY09448^,* was crossed with *trx* mutant *(trx^1^)* and males in the progeny with *mask^EY09448^/trx^1^* genotype were scored for A5 to A4 transformation. Loss of pigmentation in A5 segment of adult male flies (marked by asterisk) was used as readout for A5 to A4 transformation [36]. As a control, progeny from a cross between *trx^1^* and *w^1118^* was scored for the phenotype. High percentage of *mask^EY09448^/trx^1^* male flies exhibited A5 to A4 transformation as compared to control heterozygous *trx^1^/+* male flies. Representative image of A5 to A4 transformation phenotype in *mask^EY09448^/trx^1^* male flies as compared to the wild type are shown. All crosses were performed in triplicates and independent t-test was applied for analysis (ns; non-significant, *; p ≤ 0.05, **; p ≤ 0.01). **(C)** Depletion of Mask results in drastic reduction in *en* expression. As a control, wing imaginal discs from progeny of *w^1118^* flies crossed with *en-GAL4,UAS-GFP* show expression of GFP overlap with en expression. As compared to the control, wing imaginal disc from the progeny of *UAS-mask^RNAi^* fly crossed with *en-GAL4,UAS-GFP* show drastically reduced level of en along with diminished levels of GFP. This indicates effect of *mask* knockdown on endogenous (anti-engrailed staining) as well as on *en-GAL4* driven GFP.

## Discussion

Our results demonstrate that *Drosophila* Mask exhibits clear antagonism normally portrayed by the trxG proteins [14,22] and Mask plays an important role in maintenance of gene activation. Genome-wide binding profile of Mask has revealed key evidence about how Mask may contribute to the process of cell fate maintenance and epigenetic cell memory. Presence of Mask at *PREs* and other known Trx and Pc binding sites within BX-C and ANT-C regions highlights the possibility of close interactions between Mask and trxG/PcG system. Moreover, association of Mask near TSS at known trxG targets across the genome strongly supports the notion that Mask is likely required to maintain gene activation. The fact that Mask co-occupies Trx and CBP binding sites and positively contributes to the maintenance of H3K27ac levels (Figure 3 and Figure 4) illustrates a close interaction between Mask and CBP. In particular, a drastic reduction in expression of trxG targets like *pnr*, *pnt, psq* (Figure 1A) and *en* (Figure 5C) after depletion of Mask strongly supports our hypothesis that Mask plays a pivotal role in maintenance of gene activation by trxG. It is further substantiated by the fact that depletion of Mask leads to a drastic reduction in global H3K27ac levels and an increase in H3K27me3 levels in *Drosophila* and human cells. The suppression of extra sex comb phenotype by *mask* mutant highlights that Mask antagonizes PcG repression and acts as an anti-silencing factor like trxG. Notably, presence of Mask at *Scr* and other homeotic genes found in Mask ChIP-seq data may explain why *mask* mutant suppresses extra comb phenotype and enhances *trx* phenotype. However, it is not clear how Mask specifically interacts with CBP or other trxG members to counteract PcG repression and maintains gene activation. In Hippo signaling pathway, Mask acts as a transcriptional co-activator of Yorkie (Yki) by shuttling Yki from cytoplasm to nucleus, which is crucial for activation of Yki target genes [17–19]. Whether Mask plays a similar role in shuttling of CBP or other trxG members to the nucleus must be investigated to decipher how exactly Mask contributes to the process of gene activation by trxG.

Mask is a large scaffold protein which contains two multiple ankyrin repeat domains and a KH domain. These domains are also conserved in mammalian orthologues of Mask [18,38]. KH domains are commonly found in proteins involved in transcription regulation and nucleic acid binding, while ankyrin repeats are required for protein-protein interactions [39,40]. Histone methyltransferases, G9a and GLP, contain ankyrin repeats and these proteins bind to H3K9me1 and H3K9me2 marks via these domains [41]. Swi6, a homolog of the *Drosophila* heterochromatin protein 1 (HP1), is another ankyrin repeat domain containing chromatin regulator in yeast which is important for gene regulation [42,43]. The ankyrin repeat domains in Mask may facilitate a potential multivalent interaction by which Mask promotes clustering of multiple trxG epigenetic factors and contributes to genome organization of transcriptionally active regions. However, further work is required to explore possible involvement of Mask in genome organization of chromatin domains of actively transcribed regions.

The multifaceted role played by Mask during fly development can be explained by the fact that it is associated with genomic loci of various components of different cell signaling pathways (Supplementary Table1). The presence of Mask binding sites in chromatin at thirty-three different genes belonging to Wnt signaling pathway, including *en* (Supplementary Table 1) is of particular significance. The fact that *en* expression was downregulated in the absence of Mask signifies Mask binding at *PREs* near *en* promoter together with Trx/Pc. Mask is reported to be a modulator of receptor tyrosine kinase (RTK) signaling pathways, contributing to proliferation, cell survival and differentiation of photoreceptors [38]. Mask also plays a pivotal role in JAK/STAT signaling by positively regulating dimerization of its receptor, Domeless [44]. It is important to highlight that components of RTK, Wnt, JAK/STAT and Hippo signaling were enriched in our Mask ChIP-seq data (Supplementary Table1). We recently reported a histone kinase Ballchen (Ball) [45] as a positive regulator of H3K27ac in *Drosophila* [26]. Importantly, Ball is known to biochemically interact with Mask via Obscurin, a muscle sarcomere structural protein in *Drosophila* [46]. Both Ball [26] and Mask also exhibit strong correlation in chromatin binding pattern with CBP, highlighting a possible Mask-Ball-CBP nexus which may act to maintain gene activation. However, this potential involvement of Ball and Mask in regulation of gene expression through CBP is most likely different from their role in muscle development through their interactions with Obscurin. We speculate that specific signaling pathways may bring Mask and trxG together to regulate genes involved in cell fate determination, maintenance of cell viability and cell proliferation.

### Conclusion

Genome-wide binding profile of Mask has revealed a hitherto unknown role of Mask in maintenance of gene activation by trxG. Mask binds at known chromatin binding sites in trxG target genes and association of Mask with chromatin primarily correlates with presence of H3K27ac, which is a hallmark of gene activation by trxG. The fact that genome-wide binding profile of Mask shows a massive overlap with CBP binding sites corroborates the Mask/H3K27ac correlation and explains why depletion of Mask results in a drastic reduction in global H3K27ac levels. The conservation of role played by Mask in maintenance of H3K27ac in fly and mammalian cells highlights an evolutionary conserved interaction between Mask and trxG. Importantly, genetic interactions of *mask* with *Pc* and *trx* mutants strongly supports the claim that Mask counteracts repression by PcG and positively contributes to the anti-silencing effect of trxG. Moreover, positive regulation of *en* gene by Mask provides further *in vivo* validation and physiological relevance that trxG require Mask for maintenance of active chromatin. Since Mask is a downstream effector of multiple cell signaling pathways, we propose that Mask is one of the missing links which may connect cell signaling with chromatin mediated epigenetic cell memory maintained by trxG. However, further molecular and biochemical investigations are required to reveal molecular mechanism how Mask affects epigenetic cell memory and how cell signaling linked to Mask may regulate gene expression governed by trxG.

## Methodology

### Cell culture

Schneider’s *Drosophila* medium (Gibco, ThermoFisher Scientific), containing 10% fetal bovine serum (Gibco, ThermoFisher Scientific) and 1% penicillin–streptomycin (Gibco, ThermoFisher Scientific) was used to culture *Drosophila* S2 cells at 25°C. Schneider S2 cells adjusted to serum-free growth medium (Invitrogen), known as D.Mel-2 cells were cultured in Express Five SFM (Gibco, ThermoFisher Scientific) containing 20mM GlutaMAX (Gibco, ThermoFisher Scientific) and 1% penicillin–streptomycin. DMEM (ThermoFisher Scientific) medium containing 10% fetal bovine serum (Gibco, ThermoFisher Scientific) and 1% penicillin–streptomycin (Gibco, ThermoFisher Scientific) was used to culture mammalian HEK293T cells at 37°C with 5% CO_2_.

### Synthesis of dsRNA and RNAi

Templates for dsRNA synthesis were amplified by PCR from cDNA samples using primers containing T7 promoter sequence. These amplicons were used to prepare dsRNA through *in vitro* transcription using T7 Megascript kit (Ambion) following the manufacturer’s instructions. Primers for *trx* were taken from second-generation *Drosophila* dsRNA library (Heidelberg 2) [47]. Further information about these primers can be found at [http://rnai.dkfz.de]. Primer for *mask* and *LacZ* dsRNA synthesis are given in (Supplementary Table 2). For *ex-vivo* RNAi, cells were treated with 20μg of respective dsRNA in 6-well plate as described previously [15,45] and harvested after 5 days of treatment for further experiments.

### RNA isolation and relative gene expression analysis

Total RNA was isolated using TRIzol and cDNA was synthesized using Invitrogen Superscript III First-Strand Synthesis System for RT-PCR following manufacturer’s protocol. qPCR was performed to analyze the relative expression of target genes as described previously [15]. Expression of target genes in test and control samples was normalized with expression of endogenous control, Actin. Relative expression was determined by ΔΔCT method as described previously [48]. The control sample was set to value one for analyzing effect on the test sample.

### Antibody generation

His-tagged-Mask fusion protein, containing amino acids 1047 to 1171 of the full-length Mask, was purified using FPLC (<u>F</u>ast <u>P</u>rotein <u>L</u>iquid <u>C</u>hromatography) based Ni-Column (GE Healthcare) for antibody production. Mask antibody was generated by injecting rabbits with 450ug of purified His-tagged-Mask followed by two 100ug booster injections administered with 14 days gap between booster injections. Serum was collected after 2 months and antibody was confirmed by using anti-Mask serum for Western blotting (Figure 4D, Supplementary Figure 4).

### Western blotting

*Drosophila* S2 cells were harvested by centrifugation and lysed in lysis Buffer (150mM NaCl, 0.05M Tris, 1% Triton X-100) supplemented with following protease inhibitors: pepstatin (0.5 μg/ml), leupeptin (0.5 μg/ml), aprotinin (0.5 μg/ml), and phenylmethylsulfonyl fluoride (PMSF) (1 mM). After transferring the supernatant to fresh tubes, 2X reducing sample buffer was added and samples were boiled at 95°C for 5 minutes. Protein samples were resolved on Novex precast Tris-acetate 3-8% gradient gels (ThermoFisher Scientific). Proteins transfer onto nitrocellulose membranes was carried out at 140V for 2 hours. Blocking was done using 5% skimmed milk prepared in PBST (PBS containing 0.1% Tween-20). Blots were probed overnight with relevant antibodies at 4°C. HRP conjugated secondary antibodies were used at 1:10,000 dilution and blots were developed with ECL reagent (GE Healthcare).

### ChIP-seq Analysis

Chromatin immunoprecipitation (ChIP) from *Drosophila* D.Mel-2 cells was carried out using anti-Mask antibody as described previously [15]. Sequencing of purified DNA was performed using BGISEQ-500 at BGI Genomics Co., Ltd. BGI SOAPnuke filter was utilized to trim low-quality reads and adaptors [49,50]. The ChIP-seq data was analyzed using Galaxy web public server [51]. Sequencing reads were mapped to *Drosophila* genome (dm6) using Bowtie version 2 [52]. Peak calling from alignment results was performed using MACS version 2 with minimum FDR (q-value) cut-off set at 0.05 for peak detection [53]. Using ChIPseeker from Galaxy server, a list of annotated peaks was generated from peak file for further analysis [54]. Heat maps of Trx-C, Trx-N, CBP, Pc, H3K27ac and H3K27me3 ChIP-seq data were generated using deepTools [55], (bamComapre, computeMatrix and plotHeatmap). Heat maps were centered on Mask peaks with coverage of +10Kb around TSSs. MEME-ChIP - motif discovery, enrichment analysis and clustering on large nucleotide datasets (Galaxy Version 4.11.2+galaxy1) was utilized to generate Discriminative Regular Expression Motif Elicitation (DREME) output of Mask and Trx ChIP-seq profiles for de-novo motif discovery [56]. List of actively transcribed genes in S2 cells [26] was used and compared with Mask bound genes to create Venn diagram. ChIP-seq data of Trx-C, Trx-N, H3K27ac and H3K27me3 were used from GSE81795 [24], CBP ChIP-seq data was taken from GSE64464 [32], while Pc ChIP-seq data was retrieved from GSE24521 [25].

### Generation of CRISPR/Cas9 knockout of *ANKHD1* in HEK293T cells

HEK293T cells, 0.6×10^6^ per well, in a 6 well plate, were transfected with a mixture of 1ug *ANKHD1* CRISPR/Cas9 KO (Knockout) plasmid (Santa Cruz, sc-405893-KO-2) and 1ug *ANKHD1* HDR (Homology Directed Repeat) plasmid (Santa Cruz, sc-405893-HDR-2) using Lipofectamine LTX Reagent with PLUS Reagent (ThermoFisher Scientific) following manufacturer’s protocol. Untreated cells were labeled as mock. *ANKHD1* HDR plasmid confer puromycin resistance and RFP (Red Fluorescent Protein) signal for stable cells selection. After 24 hours, cells were visualized under fluorescent microscope for RFP signal. Media was removed 48 hours post-transfection, and replaced with fresh medium supplemented with 3ug/ml puromycin. Both the treated and untreated cells were passaged into 10cm^3^ cell culture dish and media was replaced with freshly prepared puromycin selection media every 2 days. After seven selections, there were no colonies in mock cells while discrete colonies in *ANKHD1* knockout cells were trypsinized and transferred to a 12 well plate and allowed to grow for further analysis.

### Fly strains and genetic analysis

Following flies were obtained from Bloomington *Drosophila* Stock Center:

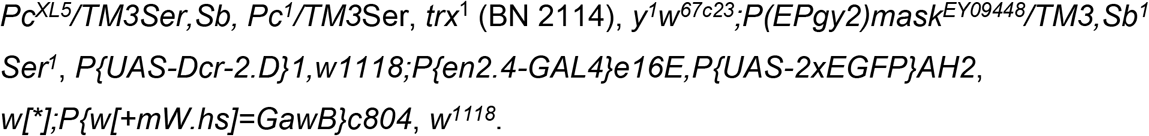

For sex comb analysis, *mask* mutant *mask^EY09448^* was crossed with *Pc* mutant alleles (*Pc^1^ and Pc^XL5^).* As a control, *w^1118^* flies were also crossed with *Pc^1^ and Pc^XL5^* for comparison. All crosses were maintained at 25°C and male progeny of these crosses were scored for extra sex comb phenotype as described previously [57]. The *mask^EY09448^* flies were crossed with *trx* mutant (*trx*^1^) and *w^1118^.* These crosses were kept at 29°C, and male F1 progeny from these crosses were scored for A5-A4 transformation phenotype. For, an *in-vivo mask* knockdown, *mask^UAS-RNAi^* fly line P{GD9632}v33396 obtained from VDRC stock center was crossed with *UAS-Dicer;en-GAL4,UAS-GFP* fly line and wing imaginal discs were immunostained with anti-engrailed antibody. For further confirmation, *mask^UAS-RNAi^* fly was crossed with a fly line (*w[*];P{w[+mW.hs]=GawB}c804*) which express GAL4 in pouch regions of wing, leg and haltere imaginal discs. Progeny from this cross was dissected at third instar larval stage and imaginal discs were stained with DAPI and anti-engrailed antibody using standard protocol as described previously [58]. For immunostaining of polytene chromosome, polytene squashes were prepared from salivary glands of third instar larvae of *w^1118^* fly. Immunostaining was performed using DAPI and anti-Mask antibody following protocol as described previously [59].

### Antibodies

Antibodies used during this study are as following: Mouse monoclonal anti-EZH2 (Sigma Aldrich, CS203195, stock concentration: 1 mg/mL diluted 1:1000 for western blotting), rabbit anti-ANKHD1 (Abcam, Ab117788, stock concentration: 1 mg/mL, diluted 1:2,000 for western blotting), mouse anti-engrailed (Developmental Studies Hybridoma Bank, AB_528224, diluted 1:50 for immunofluorescence), mouse anti-H3K27me3 (Sigma-Aldrich, Clone 18E9.1, 05-1951, diluted 1:2,000 for western blotting), rabbit anti-H3K27ac (Abcam, Ab4729, stock concentration: 1 mg/mL, diluted 1:2,000 for western blotting), mouse anti-H3 (Abcam, ab10799, stock concentration: 1 mg/mL, diluted 1:10,000 for western blotting), mouse anti-Tubulin (Abcam, ab44928, WB: 1:2000), rabbit anti-GAPDH (Abcam, Ab9485, stock concentration: 1 mg/mL, diluted 1:5,000 for western blotting), mouse anti-Lamin (Developmental Studies Hybridoma Bank, AB_528335, diluted 1:3000 for western blotting), Rabbit anti-Mask serum, ChIP: 7μL, WB: 1:250, rabbit anti-CBP serum (gift from Alexander M. Mazo, diluted 1:3,000 for western blotting).

## Acknowledgements

We would like to thank Alexander M. Mazo for anti-CBP antibody. We also thank Amir Faisal for critically reading this manuscript.

