## Supplementary Information for "Genome-wide binding of Multiple ankyrin repeats single KH domain reveals its role in maintenance of gene activation by trithorax group proteins in *Drosophila*"

**Supplementary Figure 1: Mask enrichment at *CCCT3* locus and enhancer regions mainly at *Ptx1* and *Optix* loci.**

Integrated genome browser view of ChIP-seq data highlights enrichment of Mask at enhancer regions of *Optix* **(A),** *Ptx1* locus **(B),** and highly acetylated *CCT3* locus **(C).** Mask peaks overlap with H3K27ac rich regions shown in **(C)**.

**Supplementary Figure 2: *mask* mutation results in trxG like behavior. (A)** Mutant of *mask* (*mask*$^{EY09448}$) was crossed with *Pc* mutant (*Pc*$^{XL5}$) and males in the progeny with *mask*$^{EY09448}$/*Pc*$^{XL5}$ genotype were scored for extra sex comb phenotype. Males from *Pc*$^{XL5}$ flies crossed with *w*$^{1118}$ served as a control. *Pc*$^{XL5}$/+ exhibited a strong extra sex phenotype in contrast to *mask*$^{EY09448}$/*Pc*$^{XL5}$. Total 200 male flies of the desired genotype from progeny of each cross were analyzed and based on the presence of extra sex combs on 2$^{nd}$ and 3$^{rd}$ pair of legs, flies were categorized according to the severity of phenotype. These categories are: –, no extra sex combs; +, 1–2 hairs on 2nd leg; ++, more than three hairs on 2nd leg; +++, more than 3 hairs on 2nd leg and 1–2 hairs on 3rd leg; ++++, strong sex combs on both 2nd and 3rd pairs of legs as described previously (1). The experiment was repeated three times independently and individual t-test was used to compare each phenotype category.

**Supplementary Figure 3: Mask positively regulates engrailed expression in flies.**

Depletion of Mask results in drastic reduction in engrailed (en) expression. **(A)** As a control wing imaginal disc, from the progeny of *UAS-GFP* fly crossed with *GawB-GAL4* driver line, analyzed under confocal microscope to visualize GFP signal as readout of GawB-GAL4 expression pattern. **(B)** Wing imaginal disc, from the progeny of *UAS-mask*$^{RNAi}$ fly crossed with GawB-GAL4 fly line, immunostained with anti-engrailed antibody. *w*$^{1118}$ crossed with GawB-GAL4 fly line, served as control. A drastic reduction in en expression observed as compared to control after *mask* RNAi.

**Supplementary Figure 4: Confirmation of the anti-Mask antibody generated in Rabbits.**

**(A)** Total cell lysate prepared from S2 cells was subjected to Western blotting using anti-Mask antibody. Mask protein above 400 KDa size can be clearly seen. SeeBlue™ Plus2 Pre-stained Protein Standard ladder was used for determination of protein size.

1. Tariq, M., Nussbaumer, U., Chen, Y., Beisel, C. and Paro, R. (2009) Trithorax requires Hsp90 for maintenance of active chromatin at sites of gene expression. *Proceedings of the National Academy of Sciences*, **106**, 1157-1162.
