## Supplementary figures and images for "Genome-wide binding of Multiple ankyrin repeats single KH domain reveals its role in maintenance of gene activation by trithorax group proteins in *Drosophila*"

### Supp Figure 1

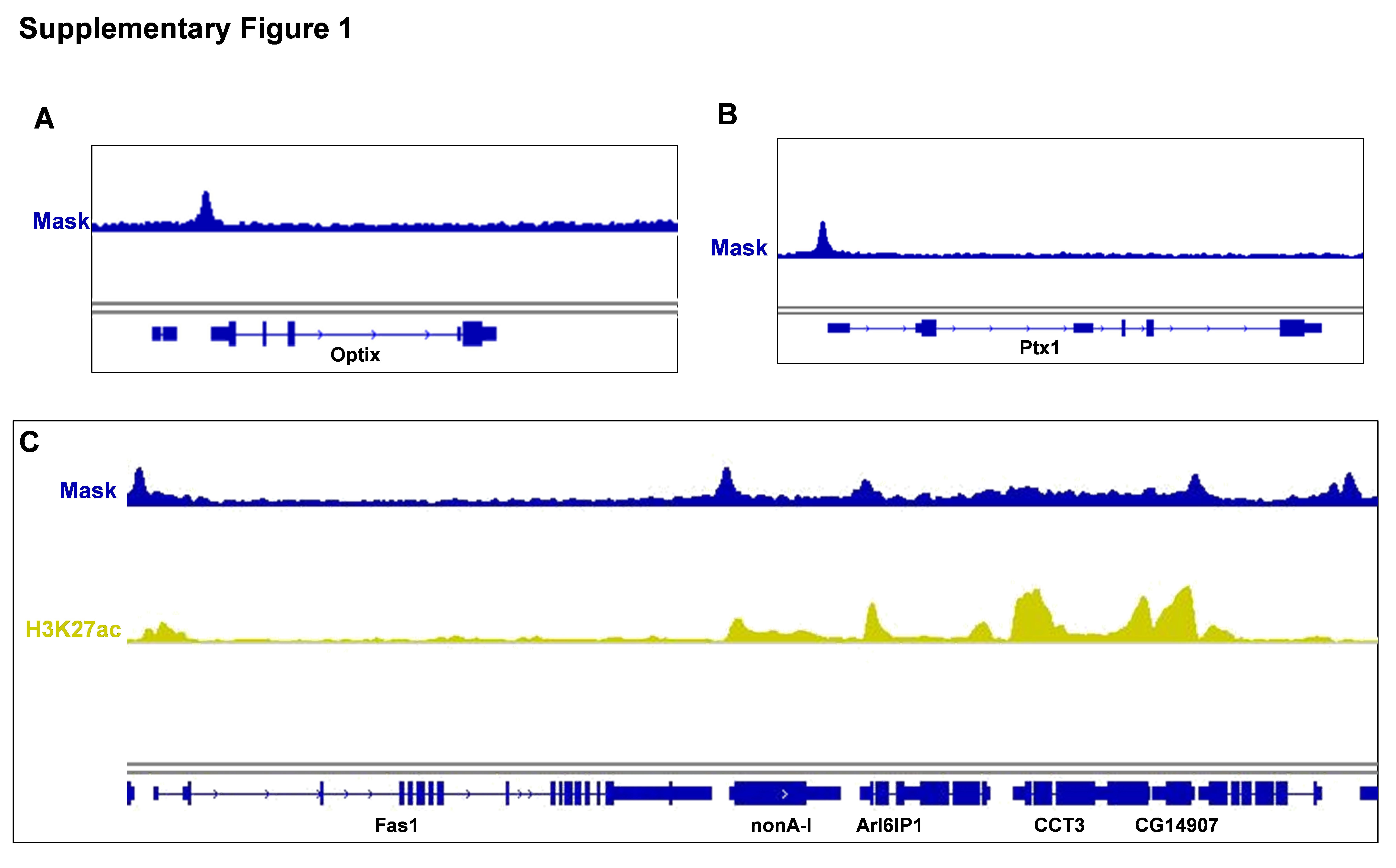

### Supp Figure 2

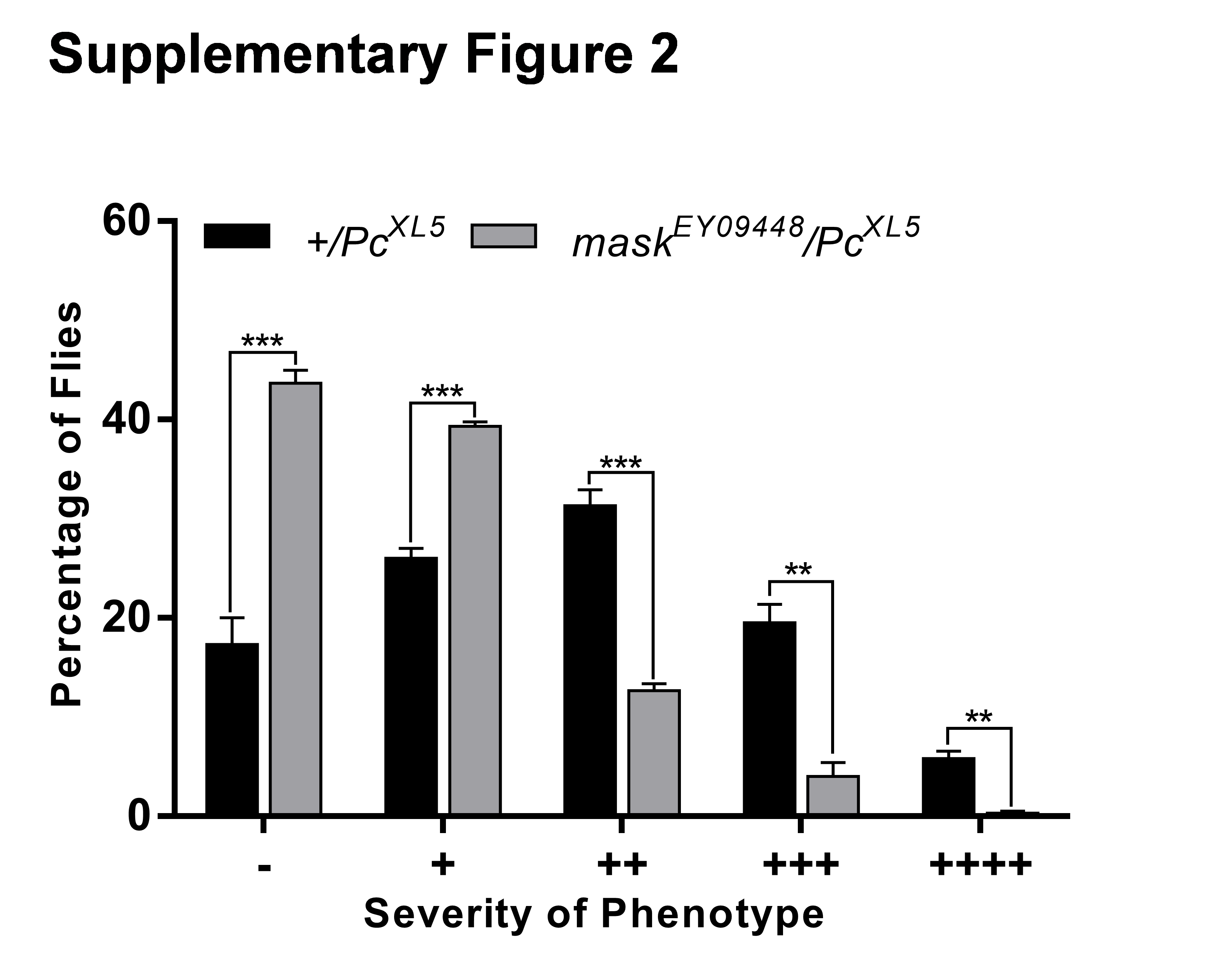

### Supp Figure 3

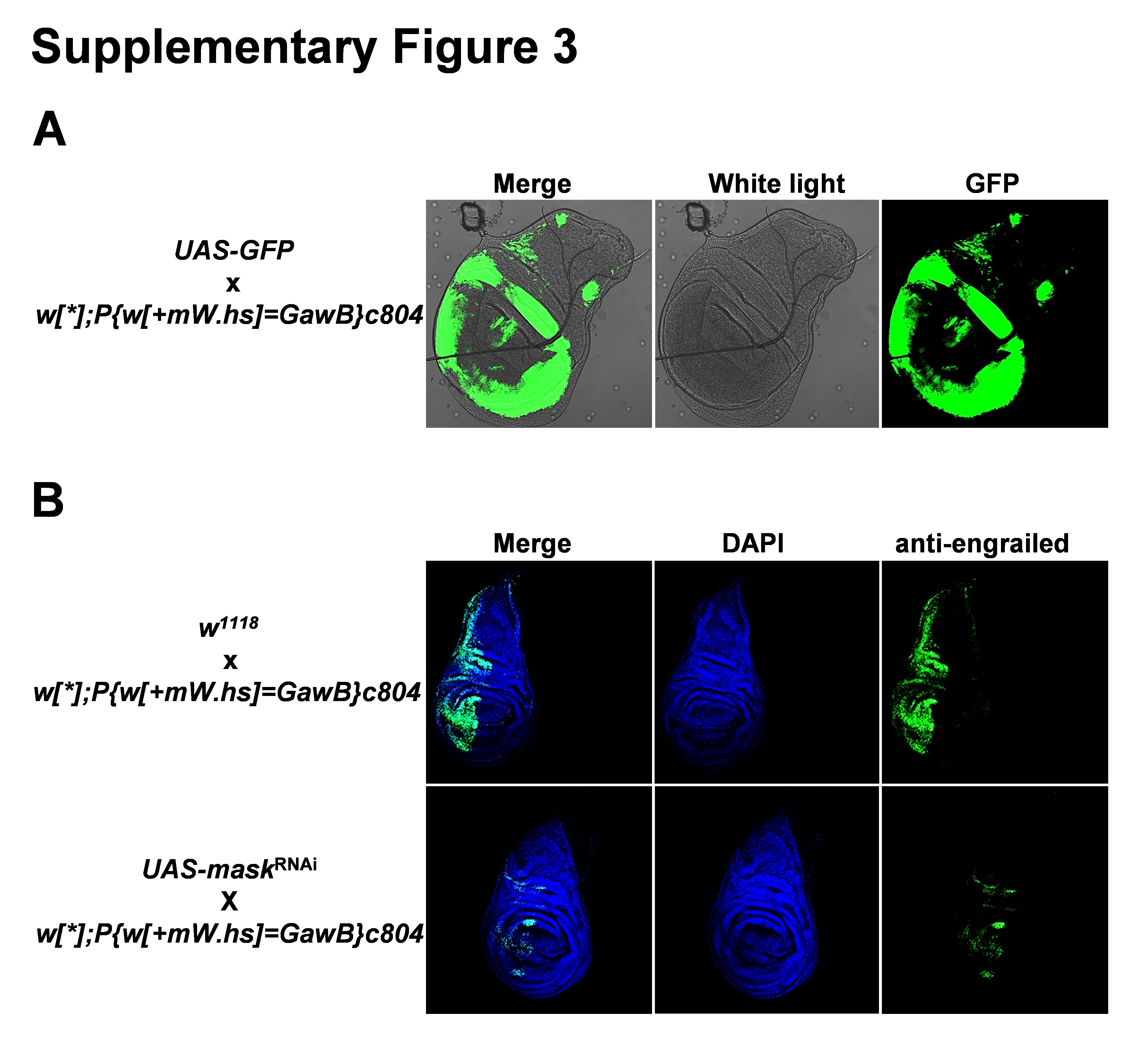

### Supp Figure 4

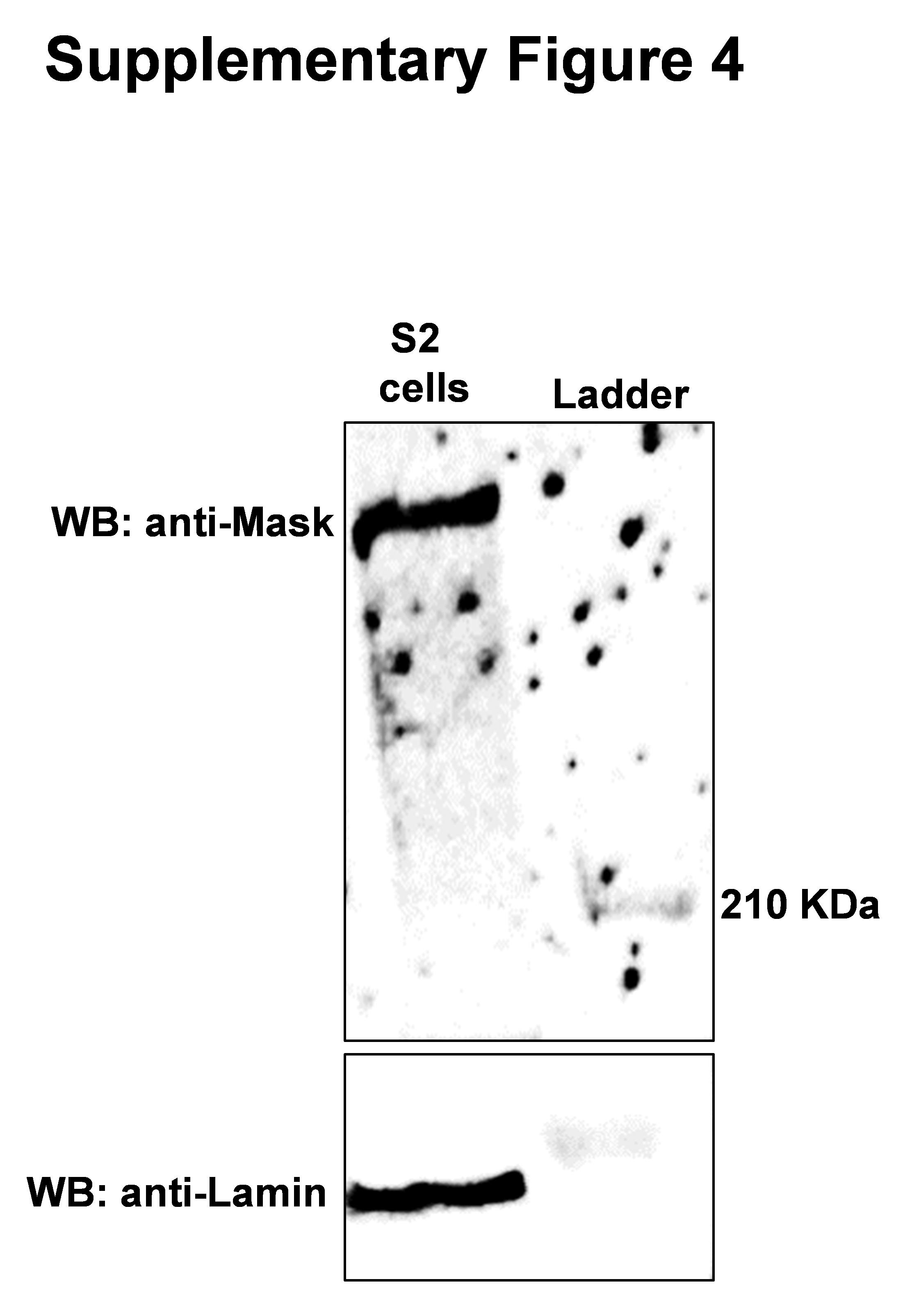
